# Real-time luminescence enables continuous drug-response analysis in adherent and suspension cell lines

**DOI:** 10.1101/2022.02.03.479010

**Authors:** Clayton M. Wandishin, Charles John Robbins, Darren R. Tyson, Leonard A. Harris, Vito Quaranta

**Author notes:** Correspondence: Clayton M. Wandishin.

## Abstract

The drug-induced proliferation (DIP) rate is a metric of *in vitro* drug response that avoids inherent biases in commonly used metrics such as 72h viability. However, DIP rate measurements rely on direct cell counting over time, a laborious task that is subject to numerous challenges, including the need to fluorescently label cells and automatically segment nuclei. Moreover, it is incredibly difficult to directly count cells and accurately measure DIP rates for cell populations in suspension. As an alternative, we use real-time luminescence measurements derived from the cellular activity of NAD(P)H oxidoreductase to efficiently estimate drug response in both adherent and suspension cell populations to a panel of known anticancer agents. For the adherent cell lines, we collect both luminescence reads and direct cell counts over time simultaneously to assess their congruency. Our results demonstrate that the proposed approach significantly speeds up data collection, avoids the need for cellular labels and image segmentation, and opens the door to significant advances in high-throughput screening of anticancer drugs.

## Introduction

Assessing cellular drug response across multiple cell lines and types is an integral component of modern cancer research. This is primarily done by taking a single cellular viability1 measurement before and after the addition of a drug across a range of concentrations in what is known as a “fixed-endpoint assay”. These measurements are then used to produce a dose-response curve to assess efficacy and potency. However, fixed endpoint assays contain a multitude of inherent biases such as the time delay effect (slow-acting drug bias), seeding density variability (T_0_), exponential growth vs. percent viability (ratio changes based on how far out the endpoint is taken), cellular growth rate dependence, and the lack of ability to produce negative values (minimum efficacy of zero) that can result in inaccurate determinations of both efficacy and potency in a variety of scenarios, potentially mischaracterizing both effective and ineffective treatments (Harris et al., 2016). A more robust alternative is to assess viability via a continuous metric. Continuous viability assays have gained substantial interest in the scientific community as they overcome the biases associated with a fixed endpoint and provide a more detailed representation of cellular drug response over time. Continuous viability assays are conducted by taking intermediate measurements across a given time interval, with short measurement intervals and extended time courses giving the most detailed information. While fixed-endpoint data yields a single number that can easily be used in dose-response curve generation, continuous assays generate multiple values, and thus require derivation to distill responses across a time period down to a single value. Assays such as EZ-MTT address this most simply by taking the slope of the dataset for dose-response curve generation, while alternative approaches such as the GR metrics and DIP rate address it by expressing each individual data series as a ratio of the basal response (Fallahi-Sichani, Honarnejad, Heiser, Gray, & Sorger, 2013; Hafner, Niepel, Chung, & Sorger, 2016; Harris et al., 2016). Continuous assays also have their own experimental hurdles that have prevented widespread adoption of the platform, such as requiring a live cell fluorescent label (direct cell counting), inefficient cell segmentation algorithms, and an inability to work well with suspension cell lines (limited by imaging ability).

Recently, a new continuous luminescence-based viability assay has been developed that indirectly measures the cellular reductive capacity through metabolic conversion of a pro-substrate to substrate (Figure 1). The novel low-toxicity and membrane permeable NanoLuc luciferase pro-substrate rapidly diffuses into cells and is converted to active substrate (Furimazine) primarily by NAD(P)H oxidoreductase, a ubiquitous and established enzyme in the cellular metabolic process (Altman, 1976; Bernas & Dobrucki, 2002; Berridge, Herst, & Tan, 2005; Berridge & Tan, 1993; Cory, Owen, Barltrop, & Cory, 1991; Duellman et al., 2015; England, Ehlerding, & Cai, 2016; Hall et al., 2012; Riss et al., 2004). Once the substrate is generated, binding to the luciferase and subsequent enzymatic cleavage produces luminescence. These luminescence values correlate well with cell counts in static measurements (Figure 2) suggesting that this system could also be used for continuous luminescence measurements as an alternative to obtaining proliferation rates by direct cell counting. This is especially promising for suspension cell cultures, where direct cell counting is often not a feasible option. Here, we show that by modifying and optimizing the commercial assay protocol for single reagent-addition, the continuous luminescence data can be used as an alternative for direct cell counting measurements. Briefly, by focusing on the rate of luminescence change in drugged cell conditions and normalizing to the basal rate of change in an undrugged population, the continuous luminescence data can be reduced to a single value, reflecting the expansion and contraction of the cell population in response to drug.

**Figure 1.**
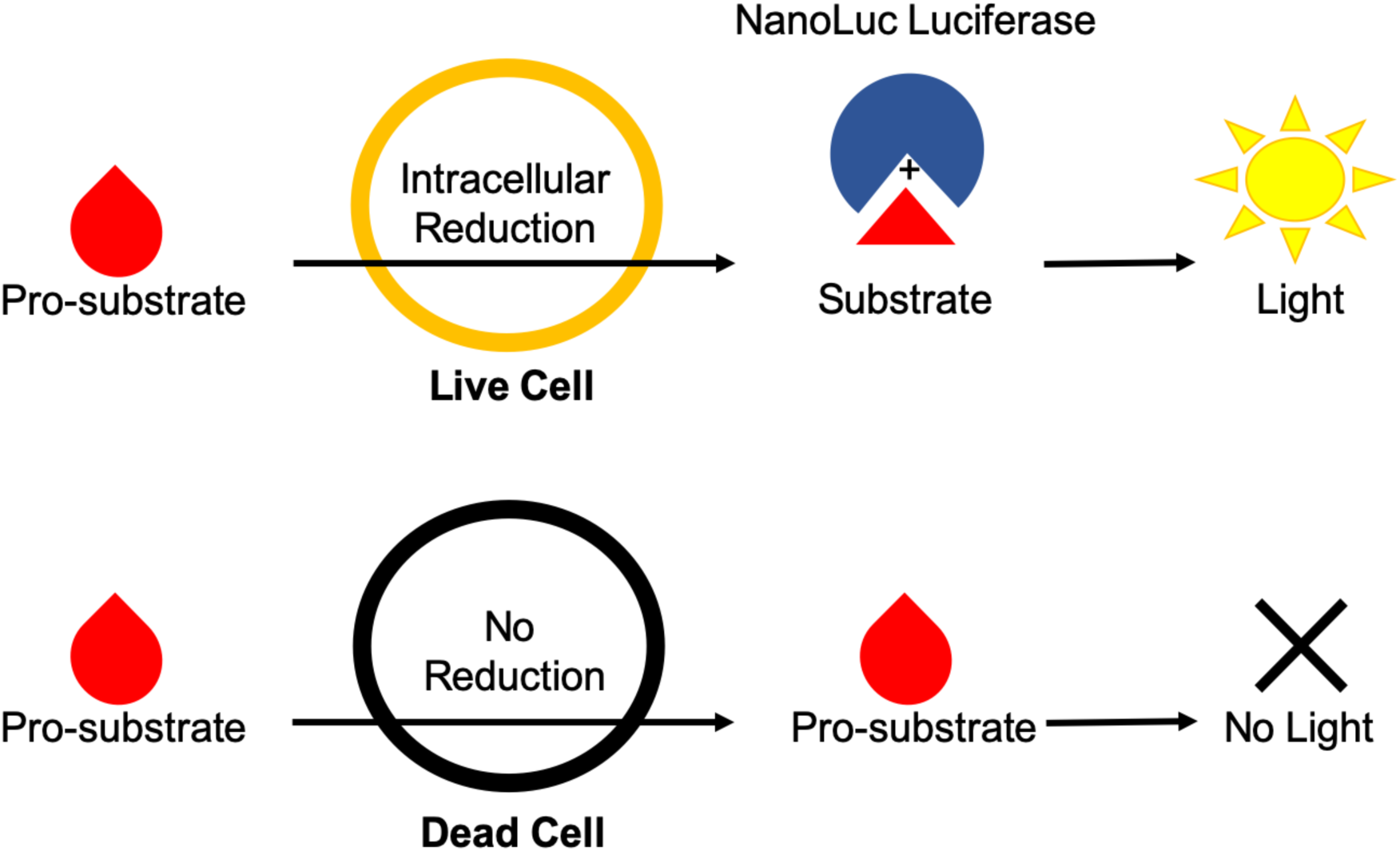
Diagram of Real-Time Luminescence Dynamics. Pro-substrate added to the culture media is rapidly metabolized by live cells via intracellular reduction into active substrate. The active substrate then reacts with NanoLuc luciferase to produce light. Dead cells are not able to metabolize the pro-substrate and therefore do not contribute to the amount of active substrate produced and subsequent light generation within the assay.

**Figure 2.**
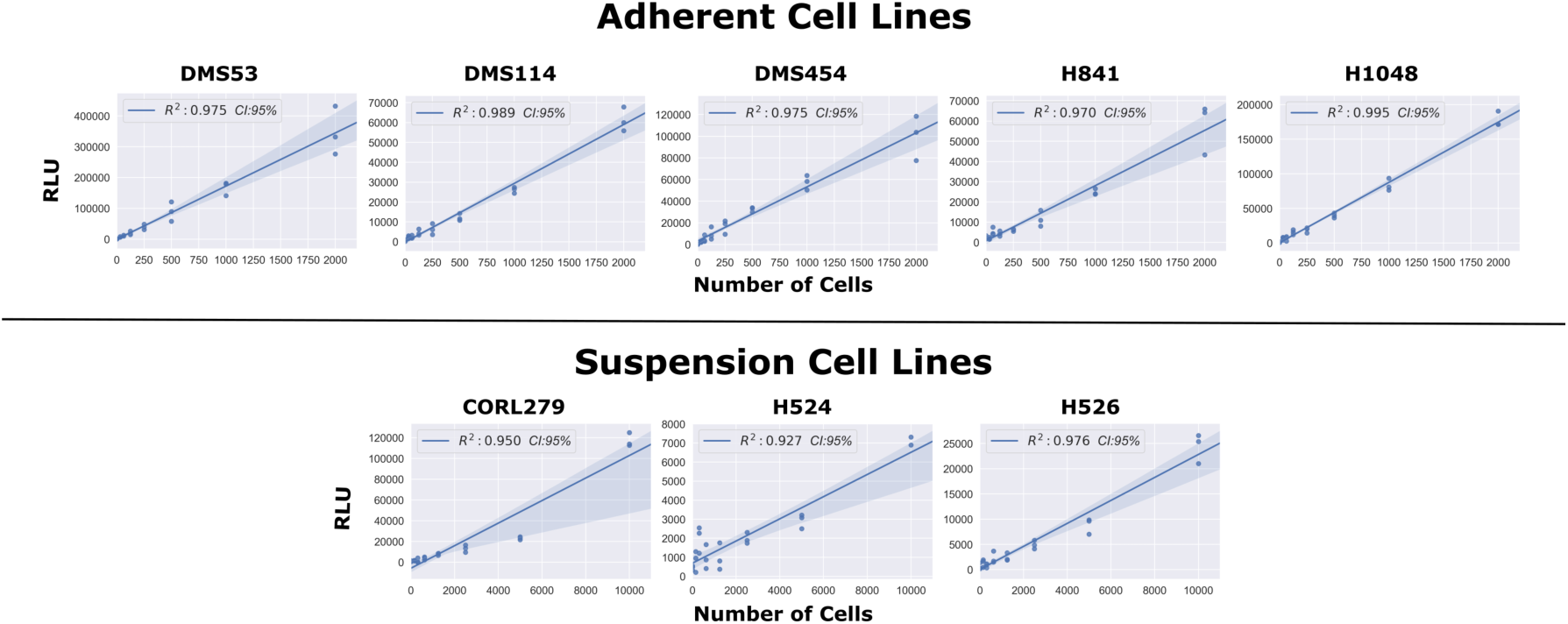
Comparison of Static Luminescent Signal and Cell Count. A range of cell lines were serially diluted by a factor of 2 from either 10,000 cells (suspension lines) or 2,000 cells (adherent lines). Assay reagents were then added to the wells and the plate was allowed to equilibrate for 1 hour. The luminescence measurements were then obtained, with the above graph showing the regression values among the static measurements of luminescence compared to varying cell seeding densities.

This streamlines the quantification of the response to the level of a fixed-endpoint assay, while remaining continuous in origin (Harris et al., 2016; Hsieh et al., 2017; Isherwood et al., 2011; Riss et al., 2004). Furthermore, we addressed challenges in the data interpretation by developing a freely available open-source analytical process (coding algorithm). Overall, using continuous luminescence to measure cellular drug response allows quantification regardless of cells being in suspension or adherent culture.

1. Cellular viability is referred to herein as the amount of live, viable, cells within a well.

## Results

### Optimizing the commercial assay for single reagent-addition continuous experiments

In order to utilize the commercial NanoLuc luciferase assay for continuous experiments, we adjusted the supplied protocol. After testing of a variety of conditions addressing NanoLuc enzyme concentration, MT substrate concentration, Solubilization temperature and duration, cell seeding density, and confluency of culture prior to experimentation (data not shown), the following tenets were obtained. First and foremost, the optimal reagent preparation was found to be 20 uL of both the NanoLuc enzyme (1000X supplied) and the MT substrate (1000X supplied) dissolved in to 25mL of culture medium supplemented with 10% FBS. We found the solubility of the MT substrate specifically, to be highly dependent on temperature.

During optimization, it was observed that the assay was more sensitive to temperature fluctuations during reads than previously anticipated. In order to address this, travel time between the plate incubator and reader was reduced to a minimum, and an additional incubation delay within a pre-warmed reader was added. The resulting optimized protocol based on these findings is available at “https://github.com/QuLab-VU/RT-Glow/tree/master/RT-Glo%20Paper”.

### Comparing luminescence to direct cell counts in proliferating cell populations

We first confirmed the relationship between luminescence signal and cell number by comparing luminescence readings and direct cell counts in cultured wells with predefined numbers of cells (Figure 2, and see Methods). To this end, we took luminescence reads across serially diluted cell concentrations after addition of assay reagents followed by one hour of equilibration. These static, single time-point measurements revealed a strong linear correlation between luminescence signal intensity and cell number in five adherent and three suspension cell lines (Figure 2). These results suggested that it is possible to monitor cell proliferation via luminescence in continuous culture over time, as a substitute for the more laborious direct cell count sampling.

To test the feasibility of continuous luminescence as an alternative for direct cell counting, we cultured multiple adherent cell lines (see Methods) and took both luminescence and direct cell counts every 4 hours for 100 hours (Figure 3). Proliferation rates were then generated by taking the slope of both the raw luminescence and log transformed direct cell counting values and compared (Figure 3). The coefficient of determination (R2) between the two proliferation rates was found to be greater than 0.92 in each.

**Figure 3.**
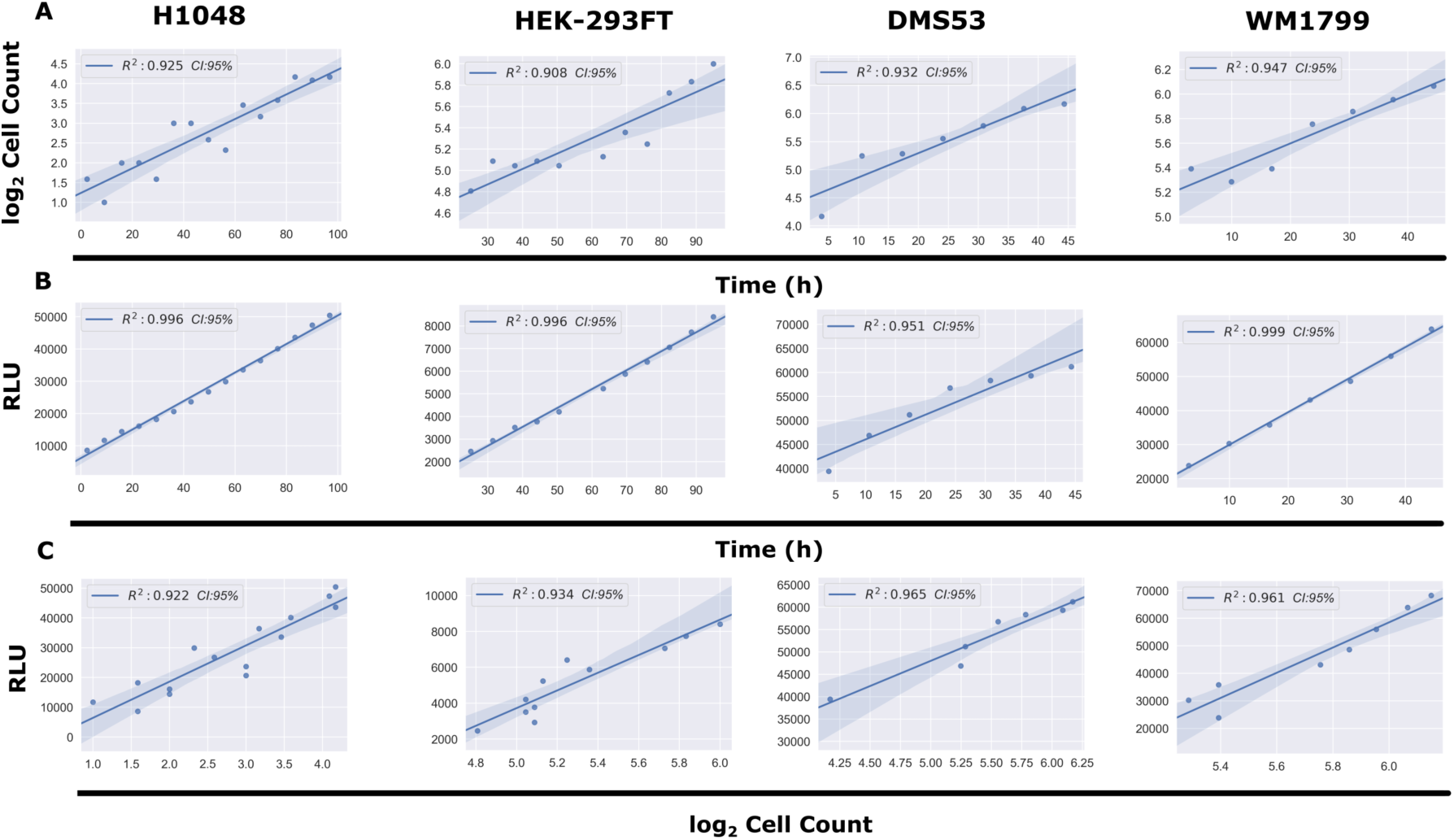
Comparison of Continuous Luminescent Signal and Cell Counts Over Time. (A) Comparison of the log_2_ transformed cell counts over time in four adherent cell lines. Cell counts were log_2_ transformed in order to linearize the data for subsequent comparisons. (B) Comparison of the continuous luminescent signal over time for the same four adherent lines from panel A. (C) Comparison of the correlation between continuous luminescent signal and log_2_ transformed cell count over time using a best-fit linear regression model. All conditions show R2 correlation coefficients >0.92.

Next, we took continuous luminescence measurements on suspension cell lines, where direct cell counting is not available, to assess if their luminescence remained linear for the duration of the experiment. Since linearity of luminescence signal is a requirement for straightforward analysis of continuous luminescence measurements (taking the slope) it was necessary to confirm this prior to using it as a metric for cell proliferation (See Methods, Determining Linear Assay Range). All three of the suspension lines tested (CORL279, H526, H1930) satisfied this requirement (Figure 4). Taken together, these results from both adherent and suspension cell cultures indicate that continuous luminescent measurements are a viable alternative to direct cell counting to assess cell proliferation over time.

**Figure 4.**
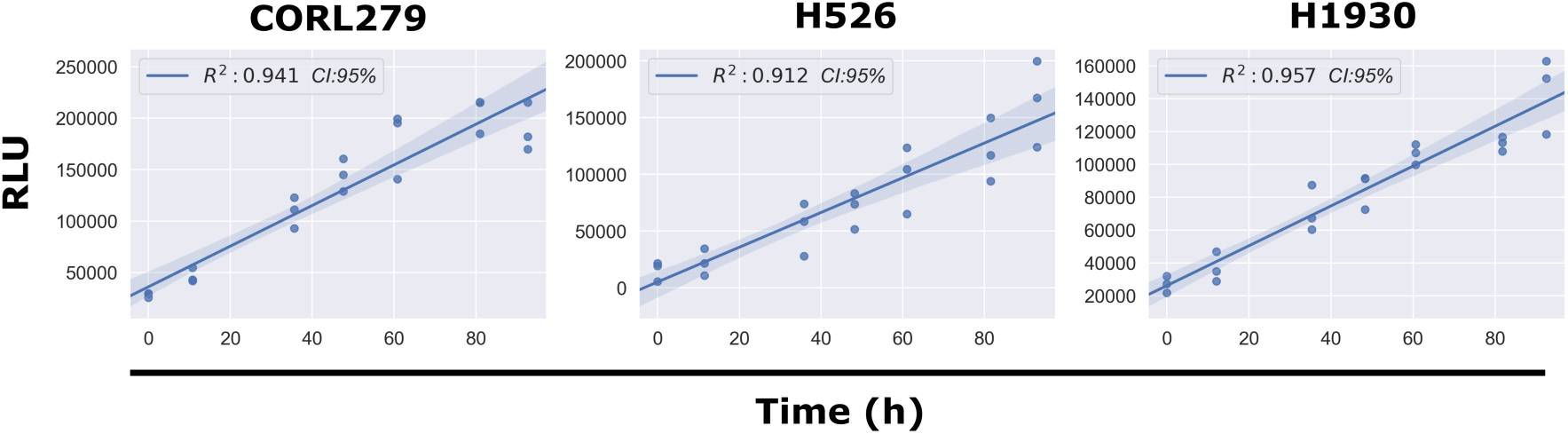
Continuous Luminescence of Suspension Cell Lines. A best-fit linear regression model of continuous luminescence in all suspension cell lines tested shows that minimum luminescent linearity requirements (R2 >0.90) are met. Real-time luminescent signal maintains a sufficient linearity for the duration of the assay.

### Quantifying drug response using continuous luminescence measurements

To explore the usefulness of the assay for continuous measurements of cell proliferation in response to drugs, we treated eight cell lines with several known anticancer agents and cultured them with the assay reagents for five days while taking luminescence measurements. Luminescence offers several advantages over conventional cell count assays (see Introduction and Discussion for more details), including speed and ease of execution and analysis for both adherent and suspension cell lines. By combining luminescence with drug-response data, continuous dose-response curves can be rapidly and efficiently generated by quantifying the rate of change in luminescence (slope). Moreover, because luminescence measurements are an indirect quantification of every single cell within a well, the data gleaned from them is much more sensitive and less variable than taking direct cell imaging counts. This is most exemplified when comparing luminescence measurements to direct cell counts produced from imaging only a fraction of a given well (standard practice).

To generate rates from the continuous luminescence data, we took the slopes of the best fit linear regression lines of the raw luminescence data. An algorithm was developed to compare increasing slices of data points from the end of the assay (defined as peak luminescence in the control condition) by calculating an R2 value for each slice, and using the highest R2 value’s linear regression slope as the basal rate for which subsequent drug dilution luminescence rates were normalized to. For drugged conditions, a similar process was used, but constrained to the region between the peak luminescence of the drugged condition, and the final timepoint of the assay determined by the peak luminescence of the control (Figure 5). Once the slopes of the continuous luminescent signals were obtained, they were normalized and plotted against the drug concentration series to obtain dose-response curves (Table 1). In comparing dose-response curves generated from luminescent or direct cell counting data; overall fitting, data variation, and EC50 values were broadly found to be in agreement (Figure 6). Across both suspension and adherent cell lines, dose-response curves from luminescence-based rates were generated successfully. The code and associated data are freely accessible in this github repository, “https://github.com/QuLab-VU/RT-Glow/tree/master/RT-Glo%20Paper”.

**Table 1.** Extracted Parameters from Best-Fit Dose-Response Models.

| Cell Line | Drug | Data Type | Hill Coef | Max Resp | EC50 | residuals |
| --- | --- | --- | --- | --- | --- | --- |
| DMS114 | barasertib | Lum | 8.754 | -0.079 | 7.86E-09 | 9.812 |
| DMS114 | AMG-900 | Lum | 0.544 | -0.234 | 1.69E-09 | 9.254 |
| DMS114 | TAK-901 | Lum | 4.181 | -0.074 | 1.6E-08 | 4.990 |
| DMS114 | YM-155 | Lum | 9.527 | -0.034 | 6.91E-08 | 13.579 |
| DMS114 | SCH-1473759 | Lum | 0.485 | -0.131 | 5.74E-09 | 7.134 |
| DMS114 | etoposide | Lum | 14.618 | -0.049 | 4.83E-07 | 16.446 |
| DMS114 | SNS-314 | Lum | 0.143 | -0.564 | 2.88E-09 | 11.205 |
| DMS454 | barasertib | Lum | 0.192 | -5.000 | 733 | 17.492 |
| DMS454 | AMG-900 | Lum | 2.816 | -1.841 | 1.17 | 23.981 |
| DMS454 | TAK-901 | Lum | 0.187 | -4.834 | 6.25E-04 | 61.902 |
| DMS454 | YM-155 | Lum | 14.273 | -0.480 | 7.6E-08 | 19.917 |
| DMS454 | SCH-1473759 | Lum | 0.490 | -5.000 | 3.81E-05 | 30.889 |
| DMS454 | etoposide | Lum | 10.439 | -1.163 | 2.99E-06 | 305.999 |
| DMS454 | SNS-314 | Lum | 0.535 | -0.511 | 1.44E-08 | 70.104 |
| CORL279 | barasertib | Lum | 0.482 | 0.143 | 5.13E-09 | 2.275 |
| CORL279 | AMG-900 | Lum | 0.047 | -0.414 | 5.44E-09 | 4.109 |
| CORL279 | TAK-901 | Lum | 1.027 | -0.114 | 4.42E-08 | 1.755 |
| CORL279 | YM-155 | Lum | 0.572 | -0.172 | 2.83E-09 | 8.129 |
| CORL279 | SCH-1473759 | Lum | 0.313 | -0.391 | 1.55E-09 | 4.560 |
| CORL279 | etoposide | Lum | 0.088 | -0.967 | 5.5E-09 | 19.581 |
| CORL279 | SNS-314 | Lum | 0.000 | -1.608 | 65.5 | 5.726 |
| H1930 | barasertib | Lum | 0.396 | 0.134 | 2.57E-08 | 4.891 |
| H1930 | AMG-900 | Lum | 0.950 | 0.500 | 7E-10 | 6.743 |
| H1930 | TAK-901 | Lum | 1.350 | 0.080 | 9.01E-08 | 6.072 |
| H1930 | YM-155 | Lum | 1.528 | -0.253 | 2E-09 | 48.869 |
| H1930 | SCH-1473759 | Lum | 0.318 | -0.550 | 5.59E-07 | 3.666 |
| H1930 | etoposide | Lum | 0.126 | -1.089 | 3.37E-09 | 7.514 |
| H526 | barasertib | Lum | 2.818 | -1.859 | 1.15 | 93.562 |
| H526 | AMG-900 | Lum | 2.821 | -1.887 | 1.11 | 144.650 |
| H526 | TAK-901 | Lum | 14.720 | 0.402 | 2.39E-06 | 23.102 |
| H526 | YM-155 | Lum | 0.299 | 0.084 | 5.12E-08 | 9.928 |
| H526 | SCH-1473759 | Lum | 13.750 | 0.101 | 2.47E-06 | 24.866 |
| H526 | SNS-314 | Lum | 0.000 | -0.103 | 2.18E-09 | 101.411 |
| H1048 | barasertib | Lum | 3.431 | -0.209 | 1.44E-08 | 10.332 |
| H1048 | hygromycin_b | Lum | 0.577 | 0.635 | 1.1E-04 | 13.097 |
| H1048 | trametinib | Lum | 1.054 | 0.238 | 1.37E-07 | 12.033 |
| H1048 | SCH-1473759 | Lum | 1.333 | -0.219 | 2.86E-09 | 17.029 |
| H1048 | YM-155 | Lum | 0.995 | -0.186 | 2.74E-09 | 4.925 |
| H1048 | TAK-901 | Lum | 2.508 | -0.209 | 1.45E-08 | 17.176 |
| H1048 | SNS-314 | Lum | 2.507 | -0.069 | 4.95E-09 | 14.051 |
| H841 | barasertib | Lum | 0.520 | -0.427 | 3.96E-09 | 44.403 |
| H841 | hygromycin_b | Lum | 2.817 | -1.851 | 1.16 | 17.366 |
| H841 | trametinib | Lum | 0.799 | -0.281 | 2.1E-08 | 17.401 |
| H841 | SCH-1473759 | Lum | 3.777 | -0.363 | 4.17E-09 | 59.188 |
| H841 | YM-155 | Lum | 0.999 | -0.794 | 9.79E-09 | 86.686 |
| H841 | TAK-901 | Lum | 2.342 | -0.380 | 1.45E-08 | 13.628 |
| H841 | SNS-314 | Lum | 2.465 | -0.377 | 8.42E-09 | 14.323 |
| DMS53 | barasertib | Lum | 2.817 | -1.855 | 1.15 | 1395.696 |
| DMS53 | hygromycin b | Lum | 2.717 | 0.857 | 5.28E-08 | 117.582 |
| DMS53 | trametinib | Lum | 1.032 | -0.659 | 1.38E-08 | 85.699 |
| DMS53 | SCH-1473759 | Lum | 0.201 | -4.430 | 2.17E-03 | 47.346 |
| DMS53 | YM-155 | Lum | 0.483 | -0.696 | 2.89E-09 | 46.906 |
| DMS53 | TAK-901 | Lum | 0.203 | -5.000 | 4.42E-03 | 45.003 |
| DMS53 | SNS-314 | Lum | 0.781 | -5.000 | 1.54E-04 | 138.641 |
| DMS53 | vemurafenib | Lum | 0.230 | -5.000 | 5.49E-03 | 43.273 |
| H1048 | SNS-314 | Direct | 0.339 | -1.673 | 1.31E-08 | 3.812 |
| H1048 | trametinib | Direct | 1.009 | 0.137 | 2.29E-08 | 9.877 |
| H1048 | SCH-1473759 | Direct | 0.601 | -1.588 | 3.32E-09 | 2.783 |
| H1048 | YM-155 | Direct | 0.730 | -1.703 | 3.01E-10 | 22.686 |
| H1048 | TAK-901 | Direct | 1.692 | -1.294 | 2.11E-08 | 1.317 |
| H1048 | barasertib | Direct | 0.322 | -2.742 | 6.29E-07 | 6.424 |
| H841 | SNS-314 | Direct | 0.550 | -0.351 | 1.12E-09 | 1.208 |
| H841 | trametinib | Direct | 2.111 | 0.588 | 5.46E-07 | 0.124 |
| H841 | SCH-1473759 | Direct | 1.312 | -0.072 | 5.33E-09 | 3.166 |
| H841 | YM-155 | Direct | 1.216 | -0.818 | 2.61E-10 | 4.012 |
| H841 | TAK-901 | Direct | 2.328 | -0.633 | 6.33E-08 | 0.634 |
| H841 | barasertib | Direct | 0.442 | -5.000 | 1.55E-05 | 0.544 |

**Figure 5.**
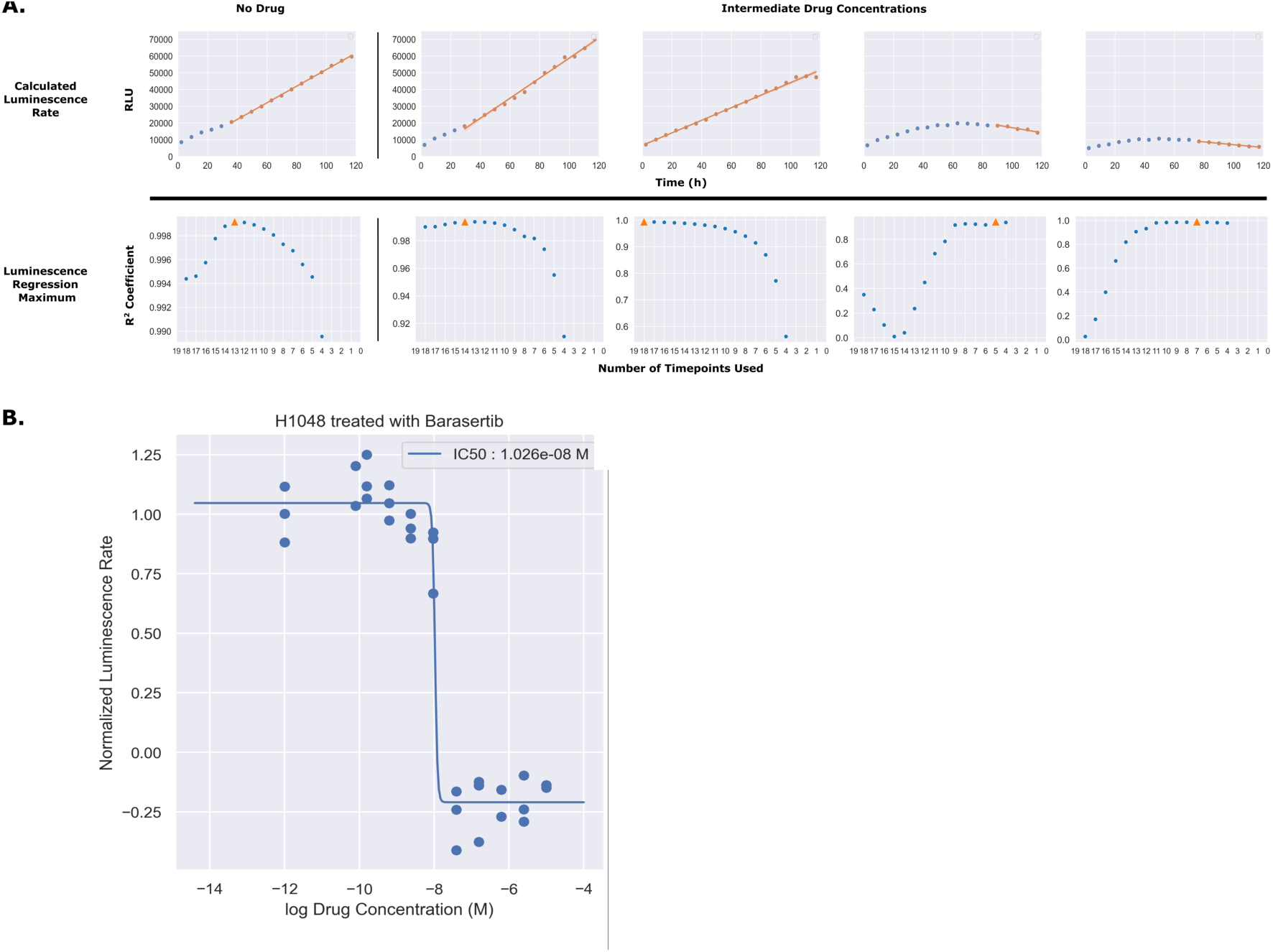
Slicing of Luminescence Data to Obtain Rate. (A) Luminescence rates for each individual drug concentration were calculated by fitting the raw luminescence data to a linear regression model. For each concentration, the number of timepoints used in the regression (slice) was determined by calculating the R2 for every possible slicing vector containing more than four points, originating from the end of the assay. The slice producing the maximum R2 value is denoted in orange as a triangle. (B) To generate dose-response curves, each of the calculated luminescence rates was normalized to the luminescence rate in the absence of drug and plotted as a normalized rate in respect to the log of the drug concentration. These data were then fitted to a four parameter log logistic function.

**Figure 6.**
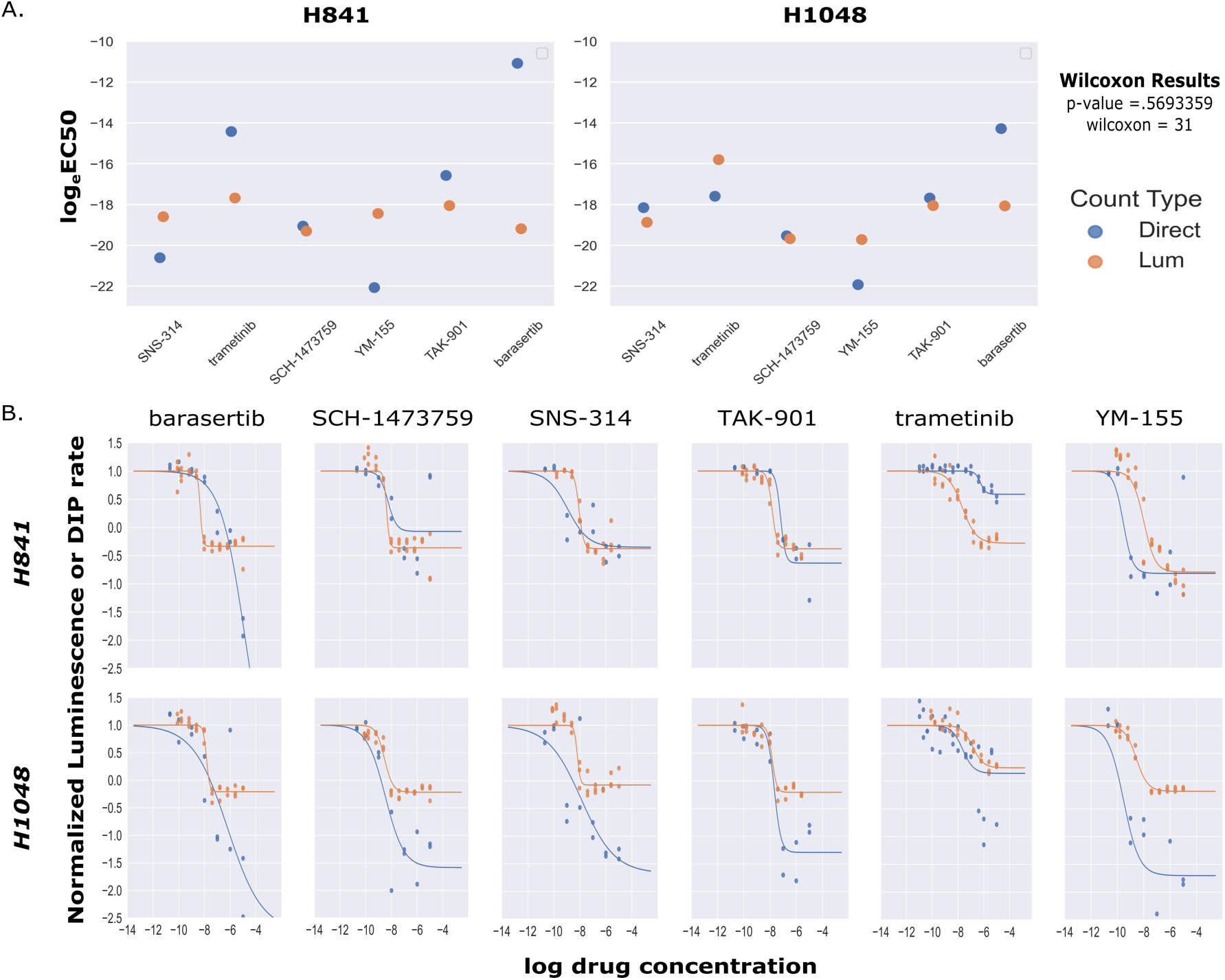
Comparison of EC50 Values and Dose-Response Curve Fits between Luminescent and Direct Cell Counting Measurements. (A) Scatter plot comparison of calculated log_e_EC50 values for both luminescence based and direct cell counting measurements. Across all paired values tested, there was no significant difference between luminescence based log_e_EC50 values and those obtained from direct cell counting (Wilcoxon Signed Rank Test, p-value=0.569, W=31, N=24). (B) Comparisons of dose-response curves generated by either luminescence (orange) or direct cell counting (blue) for two cell lines across a panel of six drugs.

## Discussion

Here we have outlined the development and application of a non-lytic luminescence-based assay to extract rate-based metrics of drug response. Implementation of our analysis and workflow has the potential to greatly expedite and modernize large-scale screening and characterization of drug response in a variety of disease models and culture methods. This work has traditionally been accomplished using fixed-endpoint viability metrics, which contain a significant degree of inherent biases, ultimately leading to a large potential for mischaracterization of drug effect in a variety of indices, both positive and negative. We and others have shown the value in taking continuous measurements across the duration of an experiment at multiple timepoints(Fallahi-Sichani et al., 2013; Frick, Paudel, Tyson, & Quaranta, 2015; Hafner et al., 2016; Harris et al., 2016; Riss et al., 2004; Uzunoglu et al., 2010). However, despite the clear advantages in data quality, adoption of continuous viability assays has been relatively slow, likely due primarily to the difficulties in integrating a continuous assay into an existing setup designed for fixed-endpoint measurements. Previously, we have described the DIP rate as an unbiased metric for drug proliferation when using direct cell counting. Our analysis of continuous luminescence utilizes the same mathematical ideology, while going one step further, with a protocol that is easily adaptable to existing fixed-endpoint workflows. What this means is, by changing only the reagent preparation method and data analysis pipeline, laboratories currently setup for drug screening using a fixed-endpoint protocol could rapidly pivot to a much more quantitatively robust method with little to no adjustment of established automation. Our hope is that this additional analytical rigor at the basic science level could lead to fewer cases of therapeutic candidates failing to translate to higher order biological models.

Like any assay, NanoLuc luciferase based continuous luminescence does have its limitations, and suffers many of the same issues surrounding MTT/MTS based measurements such as potential overestimation of viability from active mitochondrion, and inability of use for drugs targeting redox pathways(Riss et al., 2004; Wang, Henning, & Heber, 2010). These features are hardly unique to this assay, and have been generally accepted in the field for quite some time (Butcher, 2005; Cory et al., 1991; Garnett et al., 2012; Mosmann, 1983; Riss et al., 2004). By structuring experiments to avoid these known factors, complex drug-response analysis can easily be simultaneously achieved across cell lines, independent of their morphology (Wang et al., 2010).

For cell lines that are able to maintain a linear trend in luminescence for the duration of an experiment (without drug), continuous luminescence measurements offer a simple and scalable option for generating dose-response curves. This is of particular interest for cell lines that are cultured in suspension, as direct counting of suspension line cultures is not currently feasible in most situations. Based on the results of our experimentation, we intend to further explore the utility of NanoLuc luciferase based luminescence by computationally modelling the dynamics of the system, potentially using luminescence rates to predict DIP rates, as well as testing its usefulness in quantifying drug-response in three dimensional cultures (organoids). Lastly, our most immediate goal for this work is to showcase its utility with the successful integration into a high-throughput in vitro drug screening platform.

## Methods

### Cell culture

All cell lines were cultured for a minimum of two weeks prior to experimentation in T75 (Corning 430641U) flasks containing appropriate media (see below) at 37 °C and 5% CO_2_. Additionally, prior to any experimentation, absence of mycoplasma was confirmed using a MycoAlert Mycoplasma Detection Kit (Lonza LT07-118).

#### Appropriate Media

RPMI 1640 medium (Corning 10-040-CV) supplemented with 10% FBS (Gibco 26140079) and 1% Pen-Strep (Gibco 15140122)

(CORL-279, DMS53, DMS114, DMS454, H524, H526, H1048, H1930)

DMEM/F12 medium (Gibco 11320033) supplemented with 10% FBS (Gibco 26140079), and 15mM HEPES (Gibco 15630080)

(WM1799)

DMEM medium containing 4.5 g/L glucose (Gibco 11965092) supplemented with 10% FBS (Gibco 26140079), and 1% Pen-Strep (Gibco 15140122)

(HEK293FT)

### Static Luminescence measurements

Cells were cultured for two weeks, spun down, and resuspended at a density of 2.86E4 cells/mL in appropriate media, NanoLuc Enzyme (Promega E499A), and MT pro-substrate (Promega G971A). Each cell line was plated on to a 384 well GreinerOne Imaging plate (Greiner 781096) at a density of 2000 cells per well serially diluted across 10 wells (2000-4) with a total well volume of 70 uL in each. Additionally, each cell line was plated in triplicate. The plate was then incubated in a BioTek Synergy H1 at 37 °C and 5% CO_2_ for 5 minutes before luminescence measurements were taken (lid on).

### Determining Linear Assay Range

Initial cell concentrations for the linearity range of the assay were determined by following the guidelines in the “Promega RealTime-Glo MT Cell Viability Assay Protocol Handbook” under subsection four, “Determining Assay Linearity for the Endpoint or Continuous-Read Format”. Briefly, cells were serially diluted and plated with RT-Glo reagents, incubated for the proposed length of experiment (120 hours), while luminescence measurements were taken every four hours. Upon completion, the luminescence trend lines were analyzed by linear regression to find a suitable cell concentration that would maintain a linear regression coefficient of >.90 for the duration of the assay (Data Not Shown).

### Continuous Luminescence measurements

Cells were cultured for two weeks, spun down, and resuspended at a density of 4.39E3 cells/mL in appropriate media, 10 nM Sytox Green (Invitrogen S7020), NanoLuc Enzyme (Promega E499A), and MT pro-substrate (Promega G971A). Each cell line was plated on to a 384 well GreinerOne imaging plate (Greiner 781096) at a density of 300 cells per well across 10 wells with a total well volume of 70 uL in each. Additionally, each cell line was plated in triplicate. The plate was then incubated in a BioTek Synergy H1 at 37° C and 5% CO_2_ for 5 minutes before initial luminescence and fluorescence measurements were taken (lid on). The plate was then stored at 37° Celsius and 5% CO_2_ in an incubator. Every 12 hours, the plate was removed, left to equilibrate for 5 minutes in the BioTek Synergy H1, and luminescence measurements were recorded. This continued for a total of 100 hours, at which time the plates were discarded.

### Direct cell counting

To facilitate automated image processing, cells were engineered to express the monomeric red fluorescent protein mRuby2, integrated by dual transfection of a modified PiggyBac recombinase expressing plasmid and a custom mRuby2 containing transposon plasmid (Li et al., 2013; Yusa, Zhou, Li, Bradley, & Craig, 2011). Cells were seeded at 300 cells per well in 384 well GreinerOne imaging plates (Greiner 781096). DMSO (Sigma D8418) and phosphate-buffered saline (Corning 21-040-CV) were used as vehicle controls, as appropriate. Images were acquired through a 10x or 20x objective with a Cellavista HighEnd Bioimager (SynenTec Bio Services, Meunster, Germany) every 12 h as 3 × 3 or 5 × 5 montages for 120 hours. Image processing to obtain counts of cell nuclei at each time point was performed as previously described.(Frick et al., 2015)

## Acknowledgments

The authors would like to acknowledge Dr. Dave Westover for his immense help in implementing automated luminescence and imaging protocols, Dr. Jing Hao for reagent acquisition, and all members of the Quaranta and Lopez laboratories for facilitating productive discussions.

---

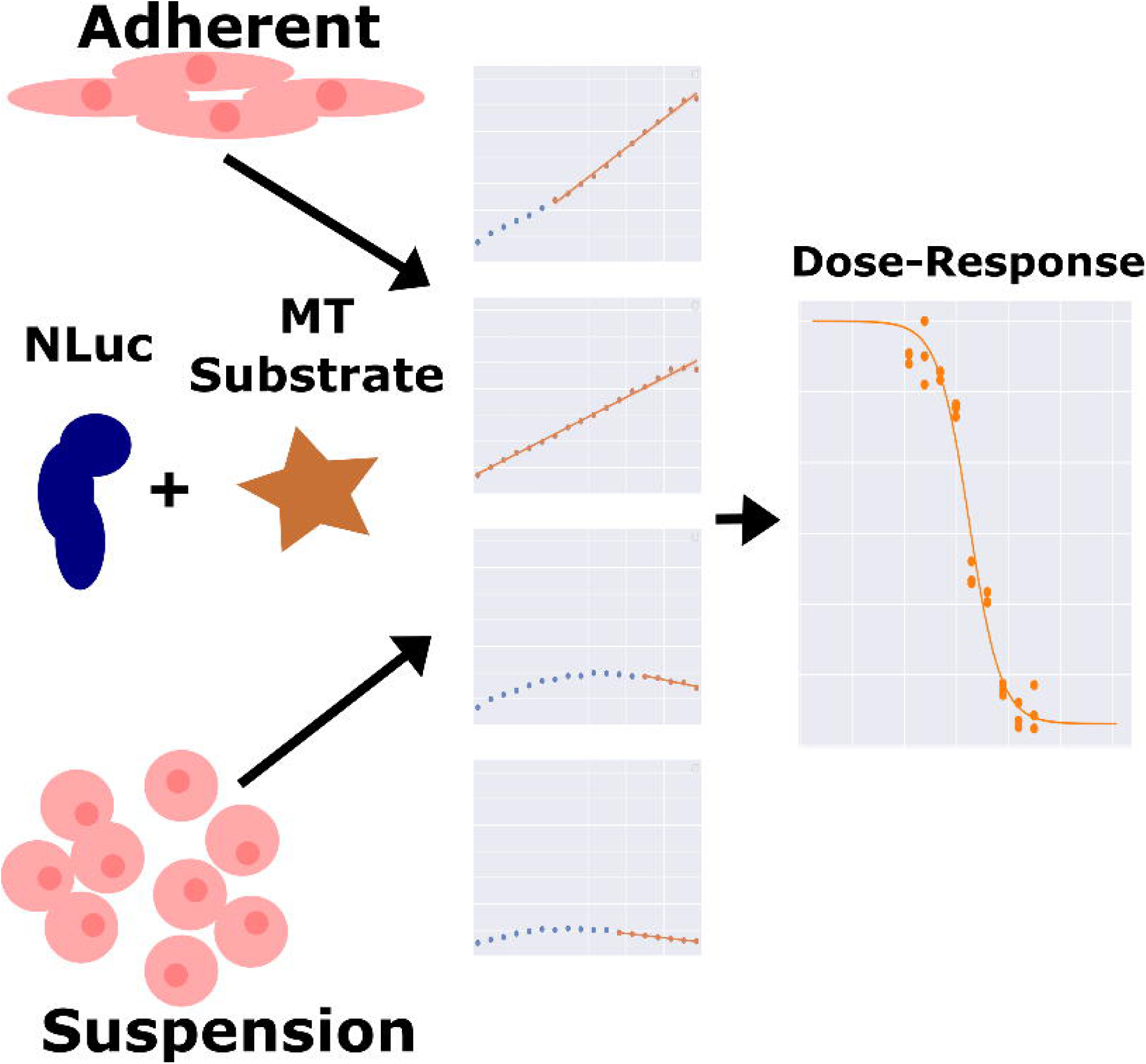

